# HES3-dependent regulatory functions in development and fusion-positive rhabdomyosarcoma

**DOI:** 10.64898/2026.05.05.723070

**Authors:** Jack Kucinski, Matthew R. Kent, Katherine M. Silvius, Ambuj Kumar, Cenny Taslim, Genevieve C. Kendall

**Affiliations:** Molecular, Cellular, and Developmental Biology Program, The Ohio State University, Columbus, OH 43210, USA; Center for Childhood Cancer Research, The Abigail Wexner Research Institute, Nationwide Children’s Hospital, Columbus, OH 43215, USA; Department of Pediatrics, The Ohio State University College of Medicine, Columbus, OH 43210, USA

**Keywords:** HES-family transcription factors, developmental biology, chromatin regulation, rhabdomyosarcoma, cancer biology

## Abstract

HES3/Her3 is a transcription factor that functions in non-canonical STAT3 signaling to promote the renewal of neural stem cells and has roles in multiple cancer contexts. To study its role in development and disease, we previously generated a CRISPR/Cas9 zebrafish knockout of *her3*, the ortholog to human HES3. HES3 is also a cooperating gene in fusion-positive rhabdomyosarcoma, an aggressive pediatric cancer, where HES3 prevents terminal myogenic differentiation, and high expression correlates with worse patient outcomes. Here, we utilize our *her3/HES3* knockout model with chromatin and transcriptional profiling techniques to assess its role during early zebrafish gastrulation with the goal of understanding the function of this transcription factor and how these activities are co-opted in cancer. We found that the Her3/HES3 preferential binding motif is distinct from other HES-family members, including a CG-rich E-box motif, that it leverages to modulate the expression of genes involved in neurogenesis and WNT signaling. We also determined that motif preferences of Her3/HES3 altered its interactions with DNA, allowing it to function canonically as a transcriptional repressor with additional duality as an activator. In the context of PAX3::FOXO1, a monogenic driver of fusion-positive rhabdomyosarcoma, we find that Her3/HES3 plays an influential role in modulating the initial activities of this core oncogenic transcription factor. Upon expressing PAX3::FOXO1 in early developing zebrafish embryos, *her3* knockout allowed for enhanced activation of neural programs, which are observed in the human disease, along with alterations to cell adhesion programs. Patient tumor samples could be clustered and stratified based on *HES3* expression alone. We saw that patient PAX3::FOXO1-positive tumors with high levels of *HES3* contained a more neural identity than those with low levels of *HES3*, altogether suggesting HES3 plays a critical role in regulating this neural signature during both the initial functions of PAX3::FOXO1 and in established tumors.

## INTRODUCTION

HES (Hairy And Enhancer Of Split) proteins are a family of basic helix-loop-helix transcription factors that function as transcriptional repressors and play important roles during embryogenesis and cell differentiation.^1^ HES transcription factors contain a DNA-binding domain, an orange domain for dimerization, and a WRPW motif for transcriptional repression, and prefer to bind to class C E-box (CACG(C/A)G) or N-box (CACNAG) motifs.^2^ While other HES-family members, like HES1 and HES5, are regulated by Notch, HES3, which we focus on, is a downstream target of STAT3, operating through non-canonical Notch signaling via the phosphorylation of serine-727 on STAT3.^3,4^ HES3 was first cloned from rat brain and embryo samples, with a human ortholog subsequently identified in silico.^5,6^ During embryogenesis, HES3 has an important regulatory role in cell fate commitment and neural differentiation as it is a downstream target of the pluripotency factor OCT4.^7,8^ Further, during gastrulation in *Xenopus laevis*, Hes3 inhibits WNT signaling to promote proper neural plate development and regulation of neural crest precursors.^9^ *HES3*, and its zebrafish ortholog, *her3*, localize to neural regions, including midbrain/hindbrain boundaries and rhombomeres.^10–12^ HES3/her3 functions to transcriptionally repress pro-neuronal targets, *ascl1* and *neurog1,* to prevent neural differentiation.^10–14^ Furthermore, HES3 helps maintain neural progenitor cells. Mice with concomitant *Hes1/Hes3* knockout present with precocious neural differentiation.^15^ *HES3* also serves as a marker for neural stem cells, and HES3 expression is downregulated following in vitro differentiation.^16^ This data suggests HES3 may be especially critical during the closely regulated processes of early embryogenesis and neural stem cell regulation.

Additionally, HES3 has a known role in various cancers. For example, in non-small cell lung cancer and glioblastoma multiforme, high levels of *HES3* expression enhances cell proliferation and viability.^17,18^ Our work focuses on the function of HES3 in fusion-positive rhabdomyosarcoma (FP-RMS), which is an aggressive pediatric cancer. The molecular drivers of this FP-RMS subtype arise from a somatically acquired chromosomal translocation, most often between *PAX3/7* and *FOXO1*.^19–21^ This translocation links the DNA binding domains of *PAX3/7* to the *FOXO1* transactivation domain, resulting in the *PAX3::FOXO1* or *PAX7::FOXO1* fusion oncogenes.^22–24^ PAX3::FOXO1 functions as a neomorphic pioneer transcription factor that can invade closed chromatin to modify the chromatin landscape and promote a tumorigenic cell identity.^25–29^ Clinically, rhabdomyosarcoma is diagnosed by the expression of myogenic markers (*MYOG, MYOD,* and *DESMA)* and is characterized by incomplete myogenic differentiation.^30^ Notably, rhabdomyosarcoma also presents in regions lacking myogenic cells.^31^ This aggressive subtype contains neural transcriptional programs and cellular populations compared to the less aggressive fusion-negative counterpart, which is instead primarily driven by RAS-mutations.^32–36^ Other HES-family members have been implicated in rhabdomyosarcoma, as depletion of HES1 and HES6 can impair tumor cell growth and tumorigenic characteristics.^37,38^ Our previous work implementing zebrafish embryonic and tumor models and cell-based systems has further highlighted a neural component of this disease. Using in vivo embryonic zebrafish models, we found that PAX3::FOXO1 initially activates neural transcriptional programs that are present in the human disease. Furthermore, we discovered that the neural transcription factor HES3 is a bona fide PAX3::FOXO1 target and cooperating gene.^28,39^ In vitro, HES3 inhibited myogenic differentiation, and its expression correlated with worse patient outcomes.^28^

Here, we aimed to investigate two questions that link the role of HES3 during developmental biology and pediatric cancer: 1) what is the function of zebrafish HES3/Her3 during early zebrafish gastrulation, and 2) how does HES3/Her3 modulate PAX3::FOXO1 functions across tumorigenic contexts? For these studies, we utilized our previously developed zebrafish *her3* knockout model.^40^ This CRISPR/Cas9 mutant contains an early termination sequence within the DNA binding domain of *her3*, resulting in a loss-of-function mutation. Through chromatin sequencing and transcriptional profiling, we identified Her3 binding preferences and subsequent developmental regulatory consequences. To further understand the functional relationship between HES3 and PAX3::FOXO1, we used a cross-species comparative oncology approach with cell culture systems, zebrafish modeling, and primary patient data to describe how HES3 modulates neural pathway activation in FP-RMS. Our work discovers new mechanisms and opportunities for HES-family member function during development and highlights avenues for HES3 to drive the more aggressive features of this pediatric cancer.

## RESULTS

### Transcription factor Her3/HES3 binds to a GC-rich class C E-box motif

We previously engineered a CRISPR/Cas9 zebrafish knockout of *her3*, the ortholog to human HES3, which resulted in a nonsense mutation within the DNA binding domain of Her3.^40^ This zebrafish mutant line is referred to as *her3^nch1^*. To understand the genomic localization and regulation of Her3 during gastrulation, we injected mRNA of either the wild-type (*her3^W^*^T^) or mutant Her3 (*her3*^nch1^) in the yolk of single-cell wild-type zebrafish embryos **(Figure 1A)**. ChIP-seq was completed at 6 hours post-fertilization, when endogenous *her3* is highly expressed **(Figure S1A)**.^41^ We identified 677 high-confidence Her3^WT^ binding sites **(Figure 1B)**. Consistent with loss-of-function activity of Her3^nch1^, there was a 3.8-fold reduction in the number of binding sites and the majority (84.6%) of Her3^WT^ sites were not detected in the mutant **(Figure S1B-C)**. Additionally, molecular dynamics simulations revealed that Her3^nch1^ resulted in a truncated protein that lacks large intrinsically disordered regions and the orange domain and contains a shortened α-helix within the DNA binding domain **(Figure S1D-G)**. Altogether, this further highlighted that Her3^nch1^ has a reduced capacity to bind to DNA.^40^

**Figure 1:**
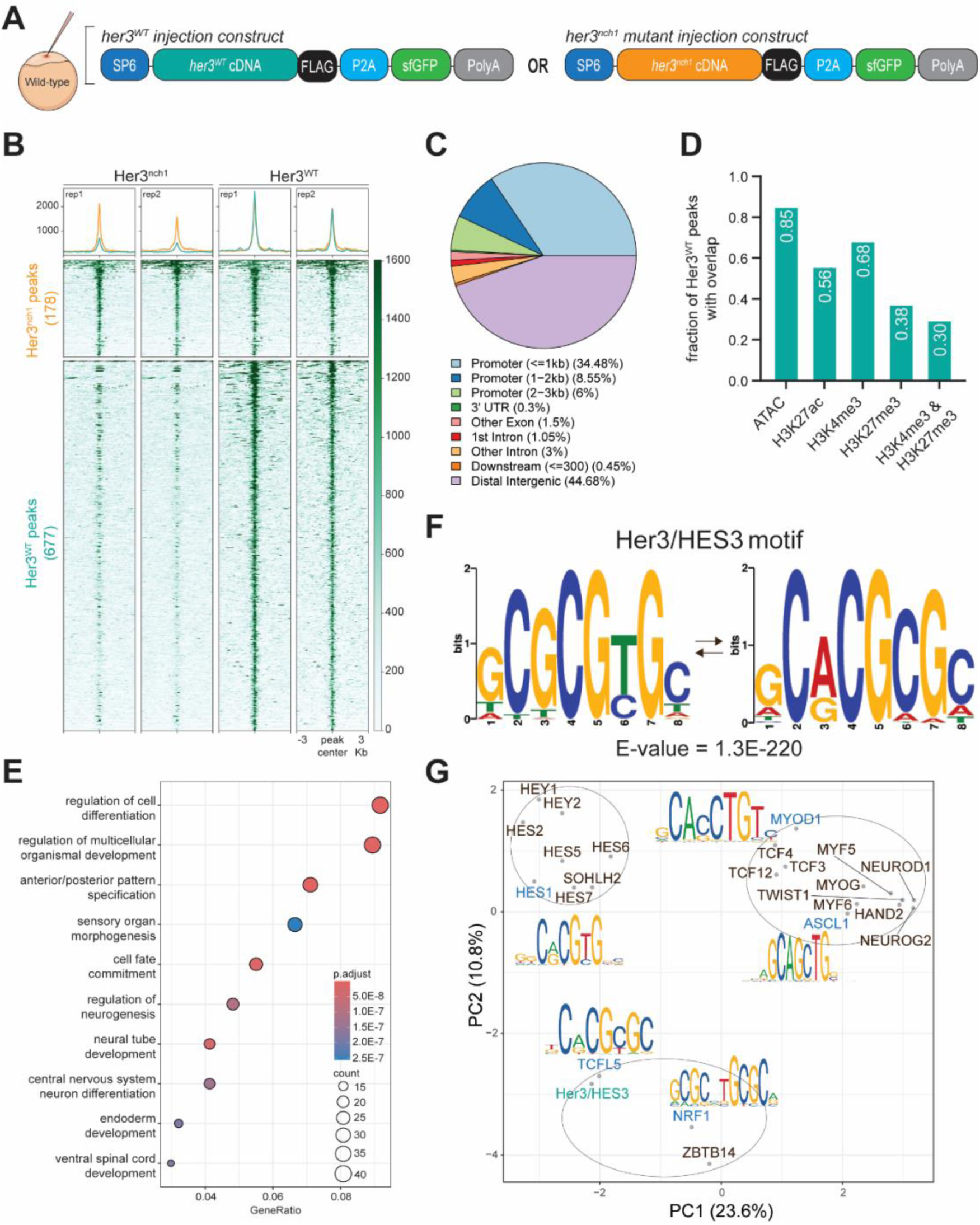
Her3/HES3 binds to a distinct E-box motif during early zebrafish gastrulation. (A) Wild-type (WT) zebrafish embryos were injected at the one-cell stage with 50 ng/μL of *her3^WT^* or mutant *her3^nch1^* mRNA. (B) ChIP-seq signal (RPKMs) at binding sites for the WT and mutant Her3 transcription factor. ChIP-seq experiments were performed in duplicate with embryos from different injection days. (C) Annotation of Her3^WT^ binding sites based on genomic localization. (D) Overlap of Her3^WT^ binding sites with chromatin accessibility and various histone marks. (E) Nearest gene pathway analysis based on Her3^WT^ binding sites. (F) The most significant motif at Her3^WT^ binding sites from MEME.^71^ (G) Principal component analysis based on motif similarity of the Her3^WT^ motif from Figure 1F to a selection of HES-family and basic-helix-loop-helix motifs.

Annotation of Her3^WT^ binding sites showed slight preferential binding to promoter sequences as 49.03% of sites were within 3 kb of an annotated gene and 68.4% of sites overlapped with an H3K4me3 peak, which marks active/poised promoters **(Figure 1C-D; S2A)**. Interestingly, there was also co-localization between Her3^WT^, H3K4me3, and H3K27me3, suggesting that Her3 may play a role in regulating bivalency during early embryonic development. Furthermore, nearest genes analysis revealed that Her3^WT^ bound near genes involved in neural and germ layer development **(Figure 1E)**. Typically, HES-family members bind to N-box (CACNAG) and E-box (CANNTG) motifs.^1^ Motif analysis at Her3^WT^ sites revealed enrichment for a distinct class C E-box motif (CACGCG) where the thymine is replaced with a cytosine **(Figure 1F)**. Similar motifs were seen with both HOMER and MEME motif analysis, but this motif preference was lost for Her3^nch1^ **(Figure S2B-C)**. This differential motif preference between HES-family members is likely driven by evolutionary divergence in the DNA binding domains as Her3/HES3 is in a specific sub-clade from other HES-family members **(Figure S2D)**.^9,42^ Luciferase reporter assays have shown that other HES-family members can bind to a similar motif in particular instances, but the high guanine/cytosine prevalence within the preferred motif appears distinct to Her3/HES3 **(Figure 1G)**.^2,43,44^ Through our ChIP-seq analysis, we discovered that Her3/HES3 binds to a distinctive GC-rich class C E-box motif.

We were intrigued to understand the relationship between DNA methylation and Her3 given the CpG dinucleotides within its binding motif. By overlapping Her3^WT^ binding with public meDIP-seq data^45^, we found that binding sites were largely devoid of DNA methylation, suggesting that methylated DNA restricts Her3 binding **(Figure S2E)**. This finding matches previous SELEX experiments, which found that HES-family members were resistant to DNA with cytosine methylation, and shows that Her3 has a similar limitation.^46^ Additionally, Her3^WT^ had limited binding to inaccessible chromatin as 85.9% of sites were in open chromatin (defined as containing an ATAC-seq peak), and had a weaker binding intensity at those few regions of closed chromatin **(Figure S2F-G)**. In summary, Her3 frequently binds to a variant GC-rich E-box motif at open promoter regions of the genome.

### Knockout of *her3/HES3* modulates broad developmental programs and WNT signaling during early embryogenesis

To understand the regulatory function of Her3 during the onset of gastrulation, we completed bulk RNA-seq in 6 hours post-fertilization *her3^nch1^* embryos for comparison to wild-type embryos. Differential expression analysis revealed 1,888 upregulated and 1,648 downregulated genes in *her3^nch1^* embryos **(Figure 2A; S3A)**. Gene over-representation analysis and gene set enrichment analysis (GSEA) revealed repression of various developmental programs and signaling pathways, like stem cell differentiation, mesenchyme development, and gastrulation, as well as upregulation of organelle fusion **(Figure 2B-C; S3B-D)**. HES/HEY-family proteins function as clock proteins during segmentation, and therefore, these findings agree with an inability of cells to properly differentiate, likely due to stem cell biology dysregulation.

**Figure 2:**
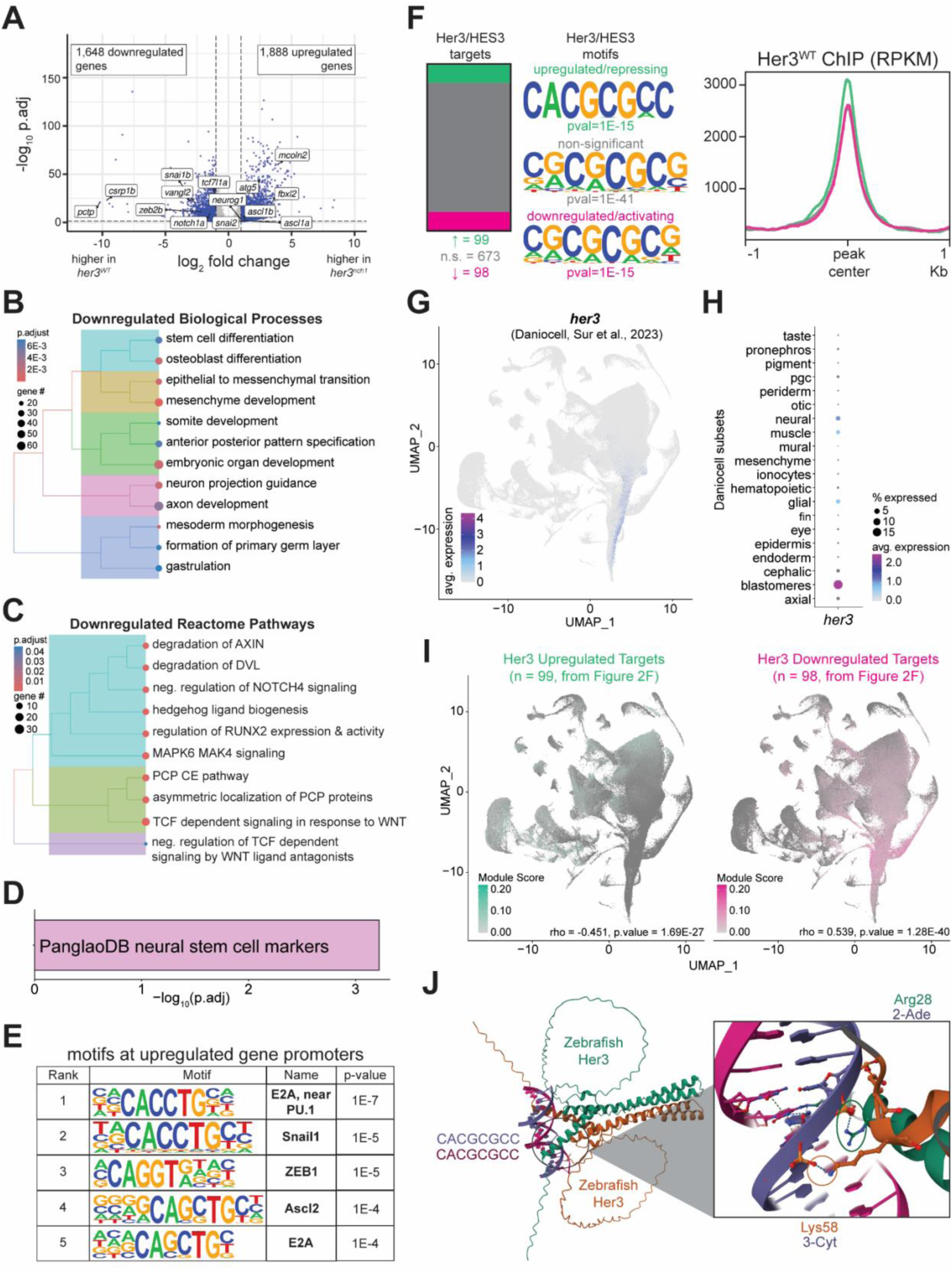
Zebrafish Her3/HES3 regulates broad developmental programs as a transcriptional activator and repressor. (A) Volcano plot of differentially expressed genes in *her3^WT^* and *her3^nch1^* embryos. Significant genes (blue) had an adjusted *p* ≤ 0.05 and an absolute fold change (FC) ≥ 2. Highlighted are manually selected genes related to neural and neural crest development. (B) Over-representation analysis (ORA) on downregulated genes for GO biological processes. (C) ORA on downregulated genes for Reactome pathways. (D) ORA on upregulated genes for PanglaoDB markers.^47^ (E) Top five known motifs at promoters of upregulated genes by HOMER.^72^ (F) Left - Fraction of Her3^WT^ targets that are differentially expressed, targets identified by combining promoter binding with the activity-by-contact model.^49^ Middle – De novo HOMER motif analysis at Her3^WT^ binding sites connected to the category of differentially expressed genes. Right – Profile plot of average Her3^WT^ ChIP-seq signal at binding sites connected to the category of differentially expressed genes. (G) UMAP of *her3* expression during zebrafish embryogenesis from Sur et al., 2023.^51^ (H) *her3* expression according to zebrafish tissue subsets from Sur et al., 2023.^51^ (I) UMAP of upregulated, left, and downregulated, right, Her3^WT^ targets from Figure 2F. Spearman correlation was completed between module score and *her3* expression across cell clusters. (J) AlphaFold modeling of Her3 binding to the motif at upregulated/repressing sites from Figure 2F.

We continued investigating WNT and neural signaling, since they are known to be influenced by HES3. During X*enopus laevis* development, Hes3 disrupts WNT signaling to promote neural plate border formation.^9^ In zebrafish, we found broad dysregulation of WNT signaling, with a greater repression of negative regulators in *her3^nch1^* embryos, suggesting HES3 modulates WNT signaling during gastrulation, in part, by an activation of its repressors **(Figure S3E-F)**. There was upregulation of known neural Her3/HES3 targets *ascl1a/b* and *neurog1* in our knockout embryos, and repression of axon development **(Figure 2A-B)**.^11,13,40^ Furthermore, we utilized cell markers from PanglaoDB to assess which cell types were potentially affected by Her3/HES3 and observed activation of a neural stem cell signature **(Figure 2D)**.^47^ Promoters of upregulated genes were also enriched for motifs for various neural and mesodermal E-box transcription factors, matching a canonical function of HES3 in keeping cells in an undifferentiated state **(Figure 2E)**. Since zebrafish neurulation does not begin until 10 hours post-fertilization, we hypothesize that Her3 functions to ensure the proper timing and regulation of these neural programs.^48^ Overall, the broad transcriptional disruption of developmental programs matches our phenotypic observation that *her3^nch1^*embryos are smaller at 24 hours post-fertilization.^40^ Altogether, HES3 plays an important role in regulating early developmental programs.

### Motif preferences and DNA interactions define Her3/HES3 functions as both a transcriptional activator and repressor

We next integrated our Her3/HES3 ChIP-seq with our previously generated ATAC-seq and H3K27ac data at this time-point to identify direct Her3 targets **(Table S1)**.^39^ Targets were identified by combining both the activity-by-contact model, which predicts enhancer-promoter interactions, and promoter-bound Her3 genes.^49^ This identified 870 Her3/HES3 targets in total, which were enriched for genes involved in pattern specification and hindbrain development **(Figure S4A)**. We saw that 11.4% (99/870) of targets were normally repressed by Her3 and upregulated in *her3^nch1^* embryos and 11.3% (98/870) targets were downregulated, suggesting they were activated by Her3 **(Figure 2F, left)**. Of note, Her3 bound near various other HES-family members, but did not result in differential expression of most of them at this early time-point **(Figure S4B)**. We further validated these targets by assessing the relationship between these genes and *her3* expression during zebrafish embryogenesis with public zebrafish single-cell RNA-seq data.^50,51^ We saw that *her3* was most highly expressed in blastomere and neural cells **(Figure 2G-H)**. Correlation analysis was completed on pseudo-bulked cell clusters. In agreement with our expectations, there was a moderate anti-correlation (rho, −0.451) between *her3* expression and genes it represses, and a moderate positive correlation (rho, 0.539) with genes it activates **(Figure 2I)**. Since HES3 is canonically viewed as a transcriptional repressor, we were intrigued by the equal distribution of directly activated and repressed genes. We excluded that Her3/HES3 transcriptional activation was a neomorphic function of the *her3^nch1^* mutation by determining that 1) Her3*^nch1^* targets were differentially expressed at the same proportion as Her3^WT^ targets, and 2) Her3^WT^ sites with significantly more binding than the mutant still had an equal fraction of downregulated targets **(Figure S4C)**. To delineate potential mechanisms for this unexplored dual regulatory role of Her3/HES3, we compared its binding characteristics between activating and repressing sites. Wherever Her3 acted canonically as a repressor, there was a higher binding intensity and enrichment for its primary class C E-box motif **(Figure 2F, middle, right)**. The non-canonical activating Her3 sites had weaker binding and enrichment for a secondary repetitive CG motif. Additionally, these sites did not have significant differences in genomic annotation **(Figure S4D)**. With AlphaFold, we modelled structural predictions of zebrafish Her3 homodimers with both motifs (CACGCGCC and CGCGCGCG), given the propensity of HES-family members to dimerize.^1^ Overall, interactions with each motif revealed similar potential binding structures, suggesting Her3 can directly bind to both motifs **(Figure 2J)**. However, investigating the second nucleotide of these motifs (A v. G) showed one dimer of Her3 would interact with the alanine of the first motif through Arg28, and Lys58 of the other dimer would interact with the third cytosine **(Figure 2J, inset)**. Meanwhile, for the guanine of the second CGCGCGCG motif, only Lys58 of one dimer was predicted to interact with that nucleotide **(Figure S4E)**. This modeling suggests that the alanine within the E-box motif allows for a second interaction between Her3 and DNA, unlike the guanine, with altered DNA interactions of Arg28 and Lys58 of Her3 homodimers. In summary, these findings and structural predictions show that differential motif preferences and binding intensities likely modify her3/HES3’s interaction with DNA and ultimately its ability to activate or repress genes.

### Overexpression of HES3 enhances oncogenic potential in fusion-positive rhabdomyosarcoma

We previously showed that high HES3 expression is indicative of worse outcomes in fusion-positive rhabdomyosarcoma (FP-RMS), an aggressive pediatric soft tissue sarcoma.^28^ Ectopic expression of HES3 inhibits terminal skeletal muscle differentiation, and HES3 is a direct target of the chimeric transcription factor PAX3::FOXO1, the most frequent driver of this disease.^28,39^ To further characterize tumorigenic mechanisms, we over-expressed HES3 in PAX3::FOXO1-positive patient-derived RH30 cells **(Figure 3A)**. We cultured cells as rhabdospheres as a proxy for transformation as 3D non-adherent spheres demonstrate enhanced chemotherapeutic resistance and xenograft tumor growth compared to cells grown in a monolayer.^52^ HES3 overexpression increased rhabdosphere formation, suggesting heightened oncogenic potential **(Figure 3B-C)**. However, transcriptional profiling following our experimental endpoint resulted in only 182 (51 up, 131 down) differentially expressed genes with HES3 overexpression. Upregulated genes were enriched for hedgehog signaling and apoptosis, while downregulated genes were involved in epithelial to mesenchymal transition (EMT) **(Figure 3D-E)**. We hypothesize that the increase in apoptosis and repression of EMT is representative of the larger sphere size when HES3 is overexpressed. Overall, these findings suggest that although HES3 does enhance oncogenicity in vitro, which matches observations in patients, its impact may not be in a tumor maintenance setting unless there is a challenging event (i.e. chemotherapy).

**Figure 3:**
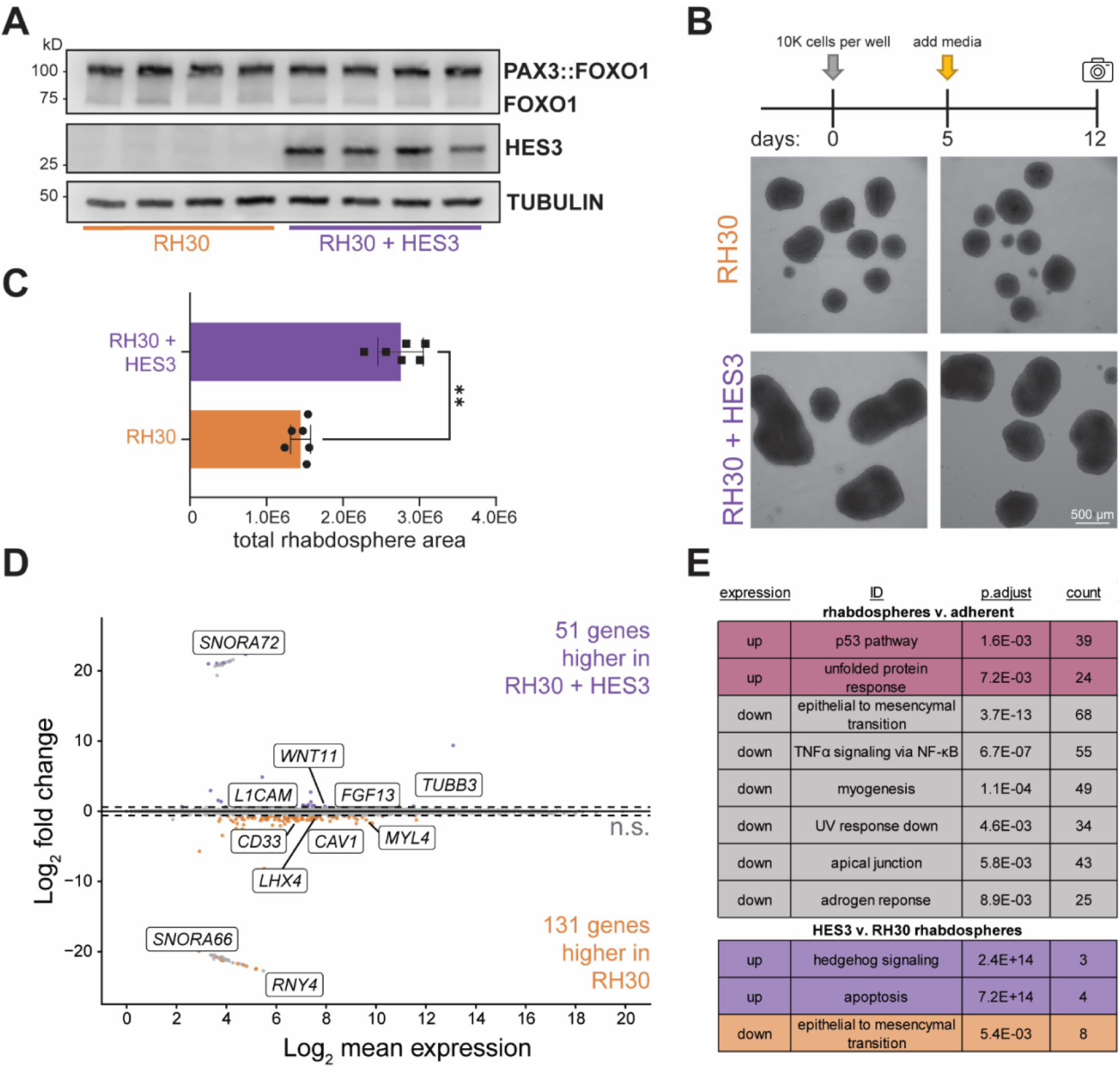
HES3 overexpression enhances fusion-positive rhabdosphere formation. (A) Representative western blot with 20 μg protein lysate from RH30 cells, a PAX3::FOXO1-positive patient-derived cell line, with or without HES3 overexpression. Each condition in the western blot contains samples from two biological replicates with two technical replicates for each. (B) Representative images of day 12 rhabdospheres. (C) Quantification of total rhabdosphere area per well. Each dot represents area of one well for 6 technical replicates per condition. A Mann-Whitney test was used to determine statistical significance. ** indicates 0.01 > *p* > 0.001. (D) Log_2_ fold change versus mean expression MA-plots for RH30 rhabdospheres with or without HES3 overexpression. Differentially expressed genes had an absolute fold change ≥ 1.5 and adjusted *p* ≤ 0.05. (E) Over-representation analysis of hallmark gene sets for Figure 3D, top, and between RH30 rhabdospheres versus adherent cell culture.

### PAX3::FOXO1 drives similar transcriptional profiles in *tp53* and *her3/HES3* deficient zebrafish backgrounds

To further decipher the regulatory mechanism of HES3 in FP-RMS, we investigated its role during the initial in vivo activities of PAX3::FOXO1. We incorporated our *her3^nch1^* zebrafish knockout line into our previously developed mRNA injection model **(Figure 4A)**.^39^ With our PAX3::FOXO1 mRNA zebrafish injection model, *her3* was upregulated and a direct target of PAX3::FOXO1 **(Figure S5A)**. In our present study, the *her3^nch1^* mutation was crossed into a *tp53*^M214K^ mutant background, which lacks normal Tp53 function, to generate homozygous double mutants.^53^ This *tp53*^M214K^ mutation was required for rhabdomyosarcoma formation in our PAX3::FOXO1 tumor model and floxing Tp53 is required in the Pax3::Foxo1 mouse model.^28,29^ We hypothesized that this *tp53^M214K^*mutation would significantly alter the initial activities of PAX3::FOXO1, since our previous work showed that misregulated Tp53 function allowed more PAX3::FOXO1-positive cells to persist (similar to the function of HES3), and *Trp53* knockouts are the most common cooperating mutation in mouse models of FP-RMS.^28,29,54^ PAX3::FOXO1 mRNA was injected into *her3^nch1^* and *tp53^M214K^* single mutants for comparison to injection into double knockouts. PAX3::FOXO1 injection into each zebrafish mutant background resulted in distinct transcriptional profiles **(Figure 4B)**. We confirmed that across all genetic backgrounds, *PAX3::FOXO1* expression levels were comparable **(Figure S5B)**. PAX3::FOXO1 still globally activated neural signatures, including markers of the FP-RMS neuronal cell population **(Figure S5C-F)**.

**Figure 4:**
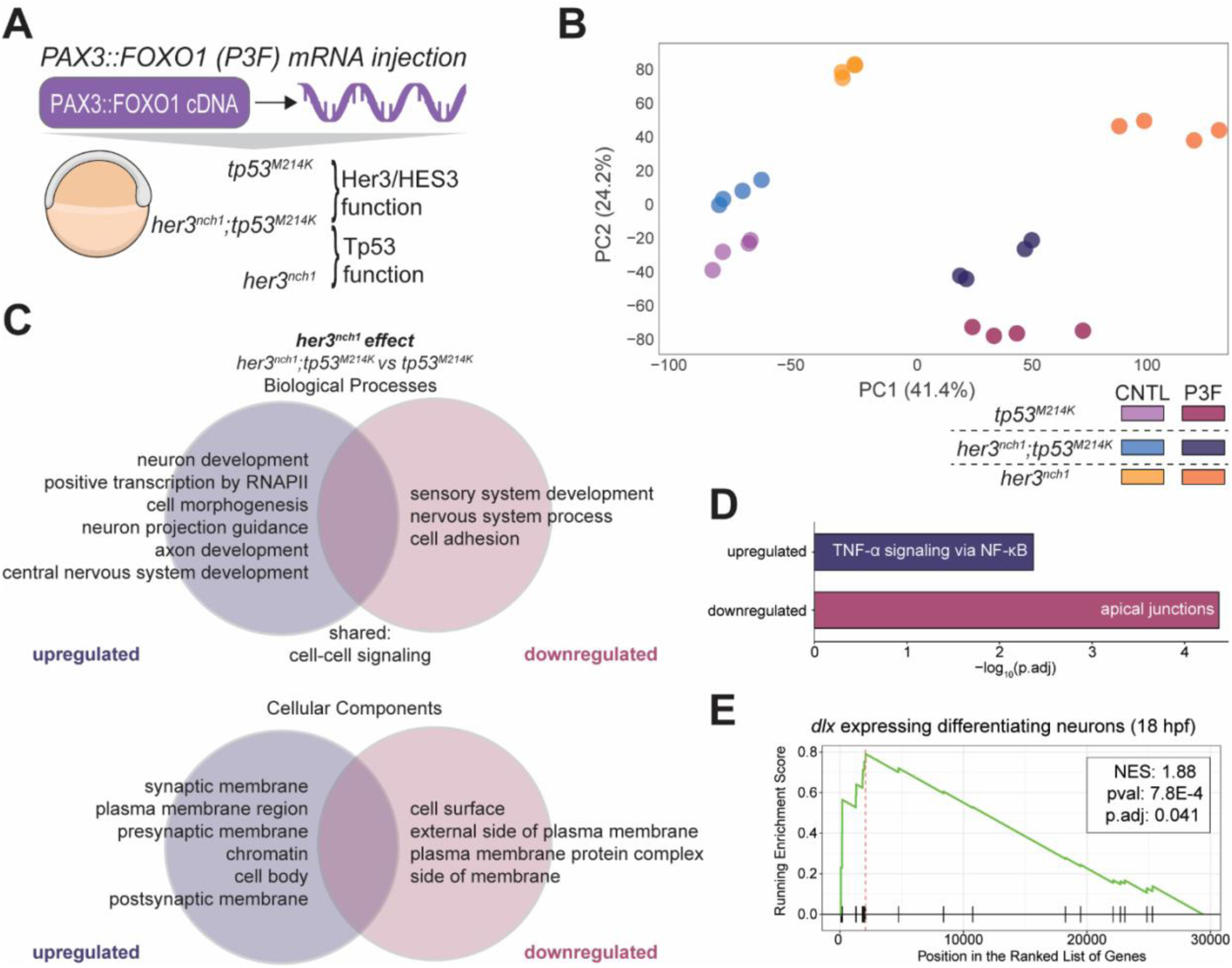
PAX3::FOXO1 has enhanced neural pathway activation in *her3* knockout zebrafish embryos. (A) Mutant zebrafish embryos were injected in the yolk at the one-cell stage with 100 ng/μL of PAX3::FOXO1 (P3F) or equal molarity of control (CNTL) mRNA. (B) Principal component analysis on RNA-seq of zebrafish mutant embryos injected with P3F or CNTL mRNA. (C) Over-representation analysis (ORA) of genes differentially upregulated with PAX3::FOXO1 injection in either *tp53^M214K^* or *her3^nch1^;tp53^M214K^*embryos. Differentially activated genes from CNTL backgrounds had an absolute fold change ≥ 2.0 and adjusted *p* ≤ 0.05. ORA to biological processes, top, and cellular components, bottom. Upregulated pathways imply they are more highly activated in *her3^nch1^;tp53^M214K^*, and downregulated pathways imply they are more highly activated in *tp53^M214K^*. (D) ORA to hallmark gene sets from genes in Figure 4C. (E) Gene set enrichment analysis of markers of the developing zebrafish cell type markers, identified from scRNA-seq analysis from Wagner et al., 2018.^89^ Comparison was made between P3F-injected in *her3^nch1^;tp53^M214K^* versus *tp53^M214K^*embryos.

We began by transcriptionally comparing *her3*^nch1^;*tp53^M214K^*and *her3^nch1^* mutant embryos to assess how Tp53 was impacting PAX3::FOXO1 functions at six hours post-fertilization, a time-point we have previously shown to have relevance to PAX3::FOXO1-driven rhabdomyosarcoma.^39^ In this early context, Tp53 supported PAX3::FOXO1’s capability to activate genes, as 1,427 genes were downregulated compared to only 434 upregulated genes when Tp53 is mutated **(Figure S6A)**. Repressed genes were involved in synaptic signaling, cell adhesion, and signaling from integrins and G protein-coupled receptors, and upregulated genes functioned in animal organ morphogenesis **(Figure S6B)**. We found that 15 of the 23 enriched downregulated pathways (p.adj = 0.01) and 88.6% (1,265/1,427) of the genes were already activated following *PAX3::FOXO1* injection in the *her3^nch1^*control background **(Figure S6C-D)**. Moreover, those genes tended to be a more highly upregulated subset of differentially expressed genes **(Figure S6E)**. Overall, it appears that Tp53 supports further activation of a particular subset of pathways that were already highly activated by PAX3::FOXO1. We hypothesized that Tp53 either directly activates these targets with PAX3::FOXO1 or indirectly allows this PAX3::FOXO1 gene signature to persist by inhibiting cell death. These possibilities are not mutually exclusive. We did not find evidence for epigenetic cooperation between these factors in our published model, as sites with increased accessibility or H3K27ac deposition upon *PAX3::FOXO1* injection largely lacked known TP53 binding motifs, and the TP53 motif was absent at the enhancers regulating these genes **(Figure S6F)**.^39^ Therefore, Tp53 is likely acting in its canonical role to prevent death, as GSEA between these PAX3::FOXO1-injected conditioned showed a decrease of an apoptotic zebrafish signature, and there was a slight rescue of the FP-RMS proliferative signature, which is typically highly repressed upon PAX3::FOXO1 injection **(Figure S6G-H)**.^39^ Taken together, this model and our analyses suggest that Tp53 indirectly supports early PAX3::FOXO1 activity by preventing cell death. This work aligns with our previous findings that *tp53^M214K^*zebrafish embryos exhibited higher numbers of PAX3::FOXO1-positive cells after injection as well as improved survival and reduced apoptosis.^28^

### Her3/HES3 knockout enhances neural pathway activation by PAX3::FOXO1

We next compared the transcriptional signatures between *her3^nch1^*;*tp53*^M214K^ double mutants and *tp53*^M214K^ mutants, which allowed us to delineate the role of Her3 in the initial functions of PAX3::FOXO1 **(Figure 4A)**. Since PAX3::FOXO1 predominantly functions as a transcriptional activator at this time-point, we focused analysis on genes that were upregulated with PAX3::FOXO1 injection in either of these two backgrounds. Of the activated genes, 706 were downregulated with *her3* knockout, i.e. higher in the *tp53*^M214K^ background and further activated by Her3/HES3 functions, and conversely, 442 genes were downregulated. Gene overrepresentation analysis revealed that removal of Her3 resulted in a repression of genes involved in cell adhesion processes and localized with plasma membrane compartments, suggesting a role in cellular migration **(Figure 4C)**. Pathway analysis to MSigDB hallmark genes revealed a similar pattern of downregulation of apical junctions **(Figure 4D)**. This is consistent with previous work where HES3 overexpression in vitro led to upregulation of *MMP3* and *MMP9*, and its developmental knockout in 24 hour post-fertilization zebrafish embryos resulted in a dysregulation of matrix metalloproteinase pathways.^28,40^ Overall, these findings further highlight a role for HES3 in cellular migration in the context of PAX3::FOXO1.

Meanwhile, PAX3::FOXO1 upregulated genes in the *her3^nch1^*;*tp53*^M214K^ double mutant compared to the *tp53*^M214K^ mutant were enriched for gene ontology terms involved in neurogenesis, such as neuron and axon development and neuron projection guidance **(Figure 4C)**. Additionally, these genes were enriched in cellular components of synaptic membranes. GSEA between PAX3::FOXO1 in both conditions agreed with a further activation of neural programs with *her3* knockout as markers for differentiated neurons became enriched **(Figure 4E)**. Lastly, analysis to hallmark genes showed an activation of TNF-α signaling **(Figure 4D)**. In FP-RMS, TNF-α-associated NF-κB has been implicated in promoting tumor cell survival, as dual pharmacological inhibition of NF-κB and BCL-2 impairs tumor cell growth.^55^ However, loss of NF-κB function does not appear to alter the rhabdomyosarcoma cell resistance to TNF-α-induced cell death.^56^ It is therefore possible that HES3 may play a role in modulating TNF-α signaling and responses to cellular stress. Overall, we observed significant differences in transcriptional changes in PAX3::FOXO1 activities, where lack of Her3 allowed further activation of neural programs, suggesting it helps to dampen and modulate this cellular program.

### *HES3* expression in patient tumors demarcates tumors with distinct transcriptional characteristics

Lastly, we investigated potential roles of HES3 in PAX3::FOXO1-positive rhabdomyosarcoma patient samples with publicly available RNA-seq data.^36^ We found that *HES3* was variably expressed between patient tumors and completely absent from skeletal muscle controls **(Figure 5A)**. Therefore, we stratified rhabdomyosarcoma tumors by *HES3* levels and compared the top quartile (HES3-high) and bottom quartile (HES3-low). Interestingly, tumor samples are well clustered based solely on the expression of this single gene, suggesting HES3 can demarcate tumors with different transcriptional characteristics **(Figure 5B)**. Differential expression analysis between groups revealed 620 genes were enriched in HES3-high tumors and 139 genes were enriched in HES3-low tumors **(Figure 5C)**. HES3-high tumors contained genes that were differentially activated with PAX3::FOXO1 injection and *her3* knockout in zebrafish, suggesting HES3 plays an active role in regulating genes in these tumors **(Figure 5D)**. Furthermore, HES3-high tumors showed a greater activation of neural biological processes, including synaptic signaling and axon development while HES3-low tumors had enriched cytoplasmic translation **(Figure 5E)**. Lastly, GSEA to markers of FP-RMS cell states showed that HES3-high tumors had a higher neuronal signature, while HES3-low tumors contained more of the proliferative and apoptotic cell states **(Figure 5F)**. Overall, this suggests that: 1) HES3 can demarcate tumors that have a greater neuronal signature, and 2) HES3 may help regulate the neuronal cell state and prevent it from shifting to proliferative and apoptotic identities.

**Figure 5:**
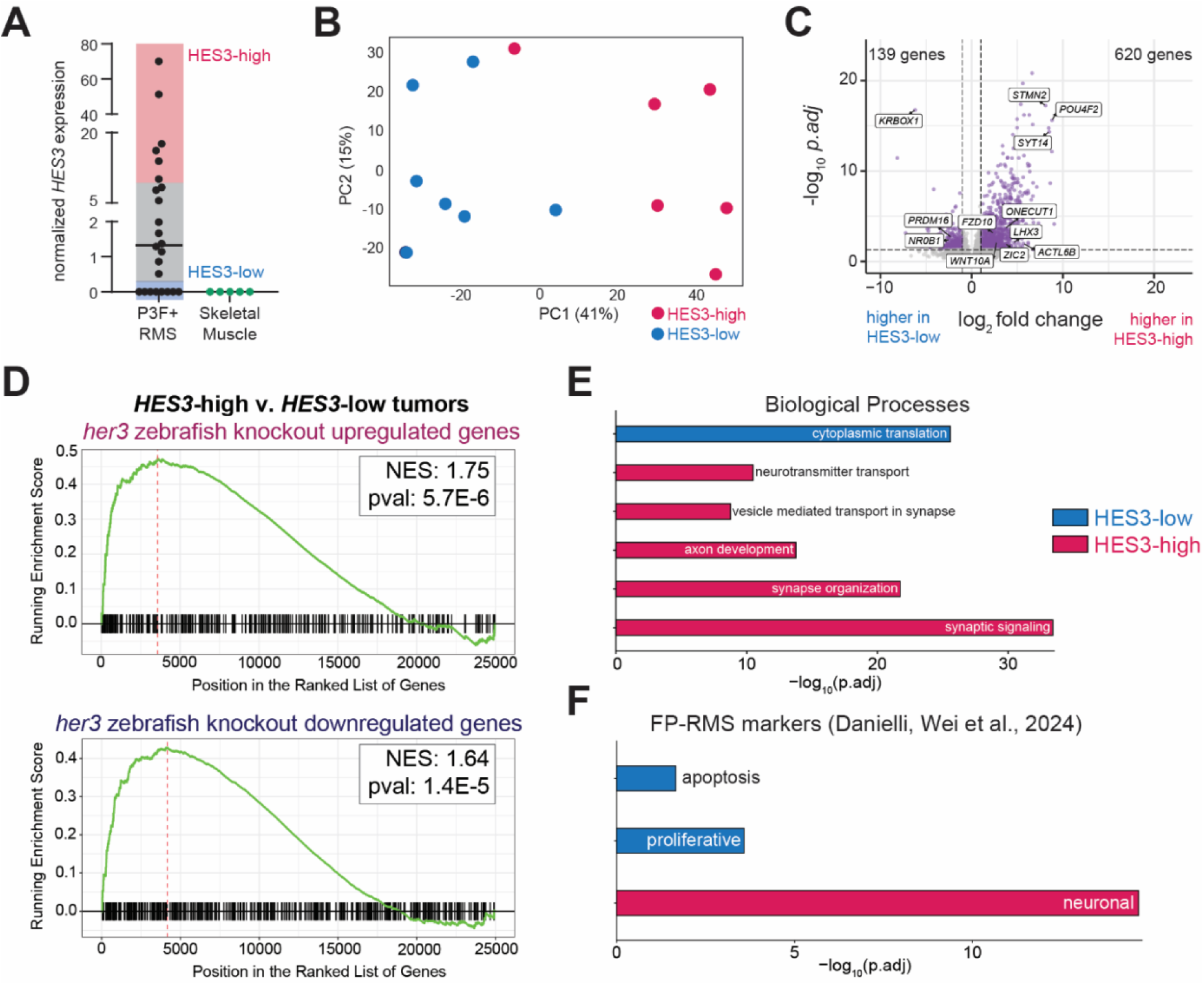
Fusion-positive rhabdomyosarcoma tumors with high *HES3* expression are enriched for neural transcriptional signatures. (A) Normalized *HES3* expression from publicly available RNA-seq in PAX3::FOXO1-positive patient tumors from Shern et al., 2014.^36^ Tumors were grouped into quartiles by *HES3* expression for comparison of top (HES3-high) and bottom quartiles (HES3-low). (B) Principal component analysis on RNA-seq of HES3-high and HES3-low tumors. (C) Volcano plot of differentially expressed genes in HES3-high and HES3-low tumors. Significant genes (blue) had an adjusted *p* ≤ 0.05 and absolute fold change (FC) ≥ 4. (D) Gene set enrichment analysis (GSEA) of genes differentially activated by PAX3::FOXO1 in zebrafish embryos with *her3* knockout from Figure 4C. GSEA to upregulated genes is on the top, and GSEA to downregulated genes is on the bottom. (E) GSEA to gene ontology biological processes between HES3-high and HES3-low tumors. (F) GSEA PAX3::FOXO1-driven rhabdomyosarcoma cell population markers identified from scRNA-seq analysis in Danielli, Wei et al., 2023 between HES3-high and HES3-low tumor.^32^

## DISCUSSION

In this work, we investigated zebrafish Her3, ortholog to human HES3, and leveraged our *her3^nch1^* knockout model to illuminate its functions during zebrafish gastrulation and to understand the initial activities of the PAX3::FOXO1 fusion-oncoprotein. During development, we found that Her3/HES3 binds to a class C E-box motif (CACGCGCC) and that *her3* loss results in broad transcriptional dysregulation of early developmental programs, especially WNT and neural signaling. In a tumorigenic context, and across various stages of fusion-positive rhabdomyosarcoma, Her3/HES3 modulates the extent to which neural pathways are present and the magnitude of neural pathway activation.

Despite being a member of the HES/HEY-family of transcriptional repressors, we found that Her3/HES3 functions as both a transcriptional activator and repressor. Her3/HES3 binding to its class C E-box motif correlates with gene repression, while binding to a repetitive CG motif is associated with activation and alters DNA interactions and binding intensity **(Figure 2F; 2J)**. Notably, other HES-family members can activate genes. By interacting with calcium/calmodulin-dependent protein kinase 2δ (CaMK2δ), HES1 activates targets during osteoarthritis development.^57^ HES6 can also interact with GATA1 to co-regulate targets, further enhancing transcriptional activation.^58^ Future studies could elucidate cooperating factors that influence HES3 activity, as seen with HES1/6. Since Her3/HES3 motifs contain CG dinucleotides and DNA methylation antagonizes Her3 binding, another potential mechanism could be passive activation through preventing the spread of DNA methylation **(Figure S2E)**. Overall, understanding the capability of HES3 and other transcription factors to switch between activating and repressing functions can provide valuable insight into context-specific and gene regulatory mechanisms.

We observed global dysregulation and downregulation of broad developmental biological processes and stem cell-related pathways in six-hour-old *her3* knockout embryos. Her3/HES3 is a downstream target of the embryonic and pluripotency transcription factor Pou5f3/Oct4.^7,8^ Our transcriptional analysis suggests that Her3 may, as a downstream effector of Pou5f3, help cells maintain an undifferentiated cellular identity instead of promoting cell fate commitment. This function does not appear solely tied to the role of Her3/HES3 in repressing neuronal differentiation and promoting stem cell renewal.^1,59^ We observed both a downregulation of axon development genes, an enrichment of neural stem cell markers, and upregulation of pro-neural factors *ascl1a/b* and *neurog1* **(Figure 2A-B; 2D)**.^12^ Our time-point of investigation occurred before zebrafish neurulation, which starts at 10 hours post-fertilization, which could explain these findings, but also highlights that Her3/HES3 helps ensure proper expression patterns and timing of neural factors even before neurulation.^48^ We also observed mis-regulation of WNT signaling in knockout embryos. HES3 interferes with WNT signaling during early *Xenopus laevis* development.^9^ In zebrafish, we find that this inhibition mainly occurs through a modulation of negative WNT regulators **(Figure 2C; S3E-F)**. In mouse embryonic stem cells, there is significant crosstalk between WNT signaling and *Stat3*.^60,61^ Given that *HES3* is a non-canonical STAT3 target, it could participate in the crosstalk and potential feedback loops between these pathways.^3^ Overall, it is clear that *her3/HES3* knockout in zebrafish causes significant transcriptional consequences during early embryogenesis and transcriptional changes that persist in one and three-day-old embryos.^40^

Our previous work found that *her3/HES3* was a primary initial target and cooperating gene of PAX3::FOXO1, a chimeric transcription factor that drives an aggressive pediatric rhabdomyosarcoma.^19,28,39^ Here, we assessed the function of Her3/HES3 during the initial activities of PAX3::FOXO1 in zebrafish and in tumor maintenance settings with cell culture systems and patient RNA-seq data. Interestingly, in 2D cell culture, HES3 does not appear to play a critical role in transcriptional regulation, which aligns with a loss of the neuronal cell population in FP-RMS in cell lines.^32^ However, even at the endpoint of rhabdosphere assays, a proxy for tumorigenic potential, there were few transcriptional differences when HES3 was overexpressed, despite HES3 driving a phenotype of larger spheres **(Figure 3)**.^52^ This finding supports that HES3 and other neural factors play a more critical role in the initial and in vivo activities of PAX3::FOXO1. Or, perhaps, the neural signatures are important after a challenging event, such as chemotherapy exposure. Previous work in fusion-positive rhabdomyosarcoma xenografts indicates these neural signatures are activated after treatment with vincristine and irinotecan.^32^ Therefore, it is possible that chemotherapy evasion is in-part an invocation of these early initiation programs. It is possible that these neural signatures, which are present in FP-RMS despite its classic association with myogenic development, may help establish or regulate the correct cellular context for PAX3::FOXO1-positive cells to persist and transform.^32–35^ Comparative analysis and lineage-tracing experiments could highlight these potential roles and mechanisms. One intriguing developmental cell population may be a transient group of *her3+/olig4+/pax3a+* cells that exist during zebrafish neural plate pre-patterning and give rise to spinal cord neural progenitors.^62^ We anticipate that research focused on the cell of origin or context-specific activities for FP-RMS could identify a neural cell population susceptible to the expression of rhabdomyosarcoma oncogenes.

In zebrafish and patient samples, we find that HES3 regulates and influences neural gene signatures. In zebrafish, *her3/HES3* knockout supports further magnitude of PAX3::FOXO1 activation of neural programs **(Figure 4C)**. Patient tumors with high *HES3* expression have transcriptionally distinct tumors with enrichment for markers of the neuronal FP-RMS cell population **(Figure 5)**. Altogether, this suggests that HES3 plays a role in preventing the hyperactivation of these pathways. In our previous work in zebrafish embryos, we found that PAX3::FOXO1 initially drives an activation of neuronal FP-RMS markers and represses proliferative FP-RMS markers.^39^ Intriguingly, the same pattern is observed during chemotherapy treatment in orthotopic mouse models and in the clustering of patient samples according to *HES3* expression.^32^ Together, these discoveries point to a balance between a proliferative or neuronal tumor identity. It will be insightful to understand the plasticity between these states and which FP-RMS population functions as a cancer stem cell. Broadly, we find that HES3 demarcates transcriptional identities and regulates neural processes. Defining the function of HES3 across tumorigenic contexts, including metastasis, recurrence, and therapeutic resistance, may provide further mechanistic insight into the aggressive characteristics of FP-RMS.

## MATERIALS AND METHODS

### RESOURCE AVAILABILITY

Further information and requests for resources/reagents should be directed to and will be fulfilled by the lead contact, Genevieve C. Kendall.

### DATA AND CODE AVAILABILITY

- All data reported in this paper will be shared by the lead contact upon request.
- New sequencing data will be deposited on GEO following revisions and acceptance of the manuscript.
- Original code to analyze the data from this paper will be posted on GitHub following revisions and acceptance of the manuscript.
- Any additional information or raw data required to reanalyze the data reported in this paper is available from the lead contact upon request.

### METHODOLOGY

#### Zebrafish Husbandry

Zebrafish (*Danio rerio)* were maintained in an AAALAC-accredited, USDA-registered, OLAW-assured, and Guide for the Care and Use of Laboratory Animals compliant facility at Nationwide Children’s Hospital. Sentinel monitoring determined that zebrafish are free of Pseudoloma neurophilia, Pleistophora hyphessobryconis, Pseudocapillaria tomentosa, Mycobacterium spp, Edwardsiella ictalurid, Ichthyophthirius multifilis, Flavobacterium columnare, and zebrafish picornavirus (ZfPV1). Adult zebrafish are housed in mixed-sex groups at a density of 8–12 fish per liter in a room with a 14:10-h light:dark cycle. A recirculating water system (Aquaneering, San Diego, CA) is used. The water is 28 °C (conductivity, 510 to 600 μS; pH, 7.3 to 7.7; hardness, 80 ppm; alkalinity, 80 ppm; dissolved oxygen, greater than 6 mg/L; ammonia, 0 ppm; nitrate, 0–0.5 ppm; and nitrite, 0 ppm) and carbon-filtered from municipal tap water, filtered through a 20-μm pleated particulate filter, and exposed to 40W UV light. Fish feed consists of live rotifer feeds for 5-30 old zebrafish, and a commercially pelleted diet twice daily for adult zebrafish. Zebrafish embryo micro-injections were performed on male and female zebrafish. WIK zebrafish were initially obtained from the Zebrafish International Resource Center (ZIRC; https://zebrafish.org/) and were used as the wild-type strain. The mutant *her3^nch1^* zebrafish used were from Kent et al., 2023.^40^ The mutant *tp53^M214K^* zebrafish line was a kind gift from Tom Look.^53^ *her3^nch1^;tp53^M214K^* double mutant zebrafish lines were established by crossing the individual mutant lines. Genotyping was completed according to Kent et al., 2023 for the her3 mutant allele and Silvius and Kendall, 2024 for the tp53 mutant allele.^40,63^ All research procedures involving zebrafish were approved by the IACUC at The Abigail Wexner Research Institute at Nationwide Children’s Hospital.

#### Zebrafish Embryo Injection

Embryos of the respective zebrafish crosses were injected into the yolk at the single-cell stage. For chromatin immunoprecipitation of FLAG tagged her3 or the her3 mutant, embryos were injected with 50 ng/μL of *her3^WT^* (Addgene: 240545) or *her3^nch1^* (pCS2+her3nch1-P2A-sfGFP) mRNA. For RNA-sequencing, embryos were injected with either 100 ng/μL of PAX3::FOXO1-2A-sfGFP (Addgene: 240098) or equal molarity of the CNTL-sfGFP (Addgene: 240100), with a drop size diameter of 0.15 mm. Injection mixes consisted of the respective mRNA and 0.05% phenol red in 3X Danieau’s buffer (52.2 mM NaCl, 0.63 mM KCl, 0.36 mM MgSO_4_·7H_2_O, 0.54 mM Ca(NO_3_)_2_·7H_2_O, 4.5 mM HEPES). The embryos were incubated in 1X E3 buffer (5 mM NaCl, 0.17 mM KCl, 0.33 mM CaCl_2_, 0.33 mM MgSO_4_) at 32°C until they reached the 6 hour post-fertilization shield stage according to Kimmel et al., 1995.^48^ mRNAs were generated via NotI digestion and SP6 reaction using the mMESSAGE mMACHINE™ SP6 Transcription Kit (ThermoFisher Scientific, AM1340).

#### Chromatin Sequencing Followed by Sequencing (ChIP-seq)

ChIP-seq was completed with *her3^WT^* or *her3^nch1^* FLAG tagged injected embryos as described in Kent et al., 2023. In brief, WIK embryos were injected at the single-cell stage with 50 ng/μL mRNA. Embryos were incubated for 5 hours and 15 minutes at 32°C to reach the developmental shield stage according to Kimmel et al., 1995.^48^ Embryos were dechorionated, deyolked, and fixed in 1% formaldehyde. Fixation was quenched with 125 mM glycine, and cells were snap-frozen for storage at −80°C. RH30 cells were prepared similarly to serve as carrier chromatin. The cell culture conditions are described below. 2 million zebrafish cells and 250,000 RH30 cells were used for immunoprecipitation and sonicated with a BioRuptor Pico in 300 μL dPBS supplemented with proteinase inhibitors. 5 μL of sonicated DNA was taken as input samples, and the remainder was incubated with 4 μg of FLAG antibody (Sigma-Aldrich, F1804; RRID:AB_262044) for 2 hours with overhead rotation at 4 °C. Incubation continued overnight with the addition of 40 μL of Protein G Dynabeads (Fisher Scientific, 10-003-D). Input and immunoprecipitated DNA were de-crosslinked with proteinase K. Library preparation and sequencing were completed as described in Kucinski et al., 2025.^39^

#### ChIP-seq Data Analysis

ChIP-seq FASTQ files were analyzed through the ENCODE ChIP-seq pipeline (https://github.com/ENCODE-DCC/chip-seq-pipeline2). Her3 peaks were determined by using the *idr.optimal* peak set following removal of blacklisted sites according to Yang et al., 2020.^64^ Bigwigs were generated by deepTools and normalized by RPKM (--binSize 1).^65^ The activity-by-contact model^49^ was used to link genes to regulatory elements from ATAC-seq and ChIP-seq data control embryos from Kucinski et al., 2025.^39^ The H3K27ac ChIP-seq data were not down-sampled for spike-in normalization. Additional publicly available sequencing data were used for this study. ATAC-seq data from Pálfy, Schulze et al., 2020 (GSE130944) and Liu, Wang et al., 2018 (GSE101779) were downloaded and analyzed through the ENCODE ATAC-seq pipeline.^66,67^ MeDIP-seq data was from Lee et al., 2015 (GSE52703).^45^ H3K4me3 and H3K27me3 data was from Akdogan-Ozdilek, Duval et al., 2022 (GSE178343).^68^ Overlap of peaks was calculated with bedTools.^69^ Principal component and differential enrichment analysis were completed with diffBind.^70^ Motif analysis was completed with MEME suite and HOMER.^71,72^ Heatmaps were generated by deepTools, v.3.3.1.^65^

#### Protein Modeling and Molecular Dynamics Simulation

Her3/HES3 protein structures were determined from the protein sequences from UniProt (Q90465 and Q5TGS1) or from the *her3^nch1^* mutant sequence.^73^ De novo structure predictions used the I-TASSER-MTD server.^74^ The top-scoring models by C-score and structural completeness were used for molecular dynamics (MD) simulations with GROMACS to refine local geometry and stabilize intrinsically disordered regions.^75^ MD simulations were performed using the CHARMM36m force field and the TIP3P explicit solvent model.^76,77^ Each model was centered in a rectangular box, solvated with water molecules, and neutralized with counterions. Energy minimization was performed using the steepest descent algorithm until the maximum force was below 1000 kJ mol⁻¹ nm⁻¹. Equilibration was conducted in two stages: 10 nano-seconds (ns) under constant volume and temperature (NVT ensemble, 300 K) using the Nose–Hoover thermostat, followed by 10 nanoseconds (ns) under constant pressure and temperature (NPT ensemble, 300 K, 1 bar) using the Parrinello-Rahman barostat with isotropic pressure coupling and a compressibility of 4.5 × 10⁻⁵ bar⁻¹.^78–80^ To enhance sampling and ensure reproducible thermodynamic behavior, three independent replicates of 200 ns production runs were performed at 303.15 K. Structural stability was assessed via root-mean-square deviation of backbone atoms and root-mean-square fluctuation using GROMACS analysis tools. Structural clustering was carried out using GROMACS to identify the most representative conformation for subsequent analysis and visualization.

Dimers of Her3 were modelled to bind to double-stranded DNA with various motifs. Motifs were flanked by four alanines to minimize edge effects. Protein-DNA interactions were modeled with alphaFold3.^81^ All protein visualization was carried out using PyMOL.^82^

#### Zebrafish RNA-sequencing (RNA-seq)

*her3^nch1^*, *tp53^M214K^*, or *her3^nch1^*;*tp53^M214K^*double mutants were injected with mRNA as described above. A minimum of 20 embryos were collected per condition for each replicate and dechorionated with pronase. Samples were snap-frozen with dry ice and stored at −80°C until RNA extraction. RNA-seq was completed in quadruplicate with injections repeated across at least two injection days to reach the desired number of replicates. RNA isolation and library generation was completed as described in Kent et al., 2023.^40^ RNA-seq analysis was completed with the nf-core/rnaseq (doi: 10.5281/zenodo.1400710) and nf-core/differentialabundance (doi: 10.5281/zenodo.7568000) with a custom danRer11 genome containing the human *PAX3::FOXO1* fusion.^83^ Differential expression cutoffs were absolute log2 fold change ≥ 1 and adjusted *p* value ≤ 0.05. Comparison of *her3^nch1^* to wild-type embryos, both injected with CNTL mRNA, utilized control embryos from Kucinski et al., 2025 (GSE270325) to mimic the stress of micro-injection from the ChIP-seq experiments.^39^ Control embryos data came from multiple sequencing runs. Pathway analysis was completed with clusterProfiler and the MSigDB database.^84–86^ Volcano plots were generated by enhancedVolcano (https://github.com/kevinblighe/EnhancedVolcano).

#### Cell Culture and Rhabdosphere Assays

RH30, a PAX3::FOXO1-positive patient-derived cell line, was obtained (ATCC, CRL-2061) and used as carrier chromatin in ChIP-seq. Additional transfected RH30 + HES3 cells (cells overexpressing HES3) or RH30 + pCMV-backbone (negative control) were from Kendall et al., 2018.^28^ Cells were authenticated by STR and tested for mycoplasma routinely through Genetica Inc. a subdivision of LabCorp. Cells were cultured in high-glucose DMEM supplemented with 10% FBS growth media and incubated in a humidified incubator at 37°C with 5% CO_2._ Transfected cells were treated with 1 ug/mL of G418. Rhabdosphere assays followed methodology and cell culture conditions from Slemmons et al., 2021.^52^ 10,000 cells were seeded per well of an ultra-low attachment 6-well plate. Cells were monitored daily and briefly swirled every two days. Images of the wells were taken during the experiment on a LEICA DMIL LED, with the total sphere area quantified by ImageJ.

Cells at the endpoint of the rhabdosphere assay and consistently cultured in normal media were isolated and snap-frozen. Total RNA was isolated for RNA-seq using an RNeasy Mini Kit. RNA library preparation, sequencing, and data analysis were completed similarly to zebrafish RNA-seq, above. For these cell lines, FASTQ files were aligned to the hg38 genome, and an absolute fold change ≥ 1.5 was used as a cutoff for differential gene expression.

#### Western Blotting

Cells were lysed in RIPA lysis buffer with protease/phosphatase inhibitors (Sigma Aldrich, R0278-500ML and Fisher Scientific, 78442). Protein concentration was determined by BCA (Fisher Scientific, PI23227). 20 µg of protein was run on a Mini-PROTEAN gel and transferred to a 0.2 μm PVDF membrane (Bio-Rad, 4568033 and 1620177). Blocking occurred with Casein and 0.05% Tween 20 (Fisher Scientific, PI37528). Primary antibodies included: 1:1000 αFOXO1 (Cell Signaling, C29H4), 1:1000 αTUBULIN (Cell Signaling, 3873S), 0.5 μg/mL HES3 (DSHB, PCRP-HES3-1A10). 1:10,000 HRP-α-rabbit (Bio-Rad, 1721019) and 1:10,000 HRP-α-mouse (Bio-Rad, 1706516) secondary antibodies were used for the corresponding primary antibody. Images were taken on a ChemiDoc Go Imaging System (Bio-Rad, 12018025) with either SuperSignal West Pico PLUS Chemiluminescent Substrate or SuperSignal West Atto Ultimate Sensitivity Chemiluminescent Substrate (Fisher Scientific, PI34577 and PIA38554).

#### Patient RNA-sequencing Analysis

Human rhabdomyosarcoma RNA-seq raw counts were obtained from Shern et al., 2014 (GSE108022).^36^ Normalization and differential expression analysis were completed with DESeq2.^87^ Samples were clustered based on top versus bottom quartile of HES3 expression. Differential expression gene cutoff was absolute log2 fold change ≥ 2 and adjusted *p* value ≤ 0.05. Samples were clustered based on top versus bottom quartile of HES3 expression.

#### Additional Statistical Analysis and Data Visualization

Additional statistical analysis was completed in GraphPad Prism 10 and R (version 4.3.0), and additional experimental information is included in the respective figure legends. Additional figures were generated with ggplot2.^88^

## ACKNOWLEDGMENTS

We are grateful for the Nationwide Children’s Hospital (NCH) Animal Resources Core for their exceptional zebrafish husbandry, especially the Zebrafish Facility team, Dr. Carmen Arsuaga, Logan Fehrenbach, Logan Bern, and Jacob Al-Armanazi. We also thank every member of the Kendall Lab for their helpful discussions and insights. Sequencing support for this project was provided by the Steve and Cindy Rasmussen Institute for Genomic Medicine and Genomics Services Laboratory. Computational support was provided by the High-Performance Computing Group at NCH for their assistance in maintaining and using the NCH cluster. G.C.K. is grateful for support from an NIH/National Cancer Institute R01 CA272872 grant, an Alex’s Lemonade Stand Foundation “A” Award, a CancerFree Kids New Idea Award, a DOD Peer Reviewed Cancer Research Program Idea Award HT94252510356, and Nationwide Children’s Hospital startup funds. J.K. and M.R.K. are supported by T32 CA269052 Training Program in Basic and Translational Pediatric Oncology Research predoctoral and postdoctoral fellowships, respectively. C.T. is thankful for support from a CancerFree Kids New Idea Award. The Institute for Genomic Medicine is funded by the Nationwide Foundation Pediatric Innovation Fund and the Ohio State University Comprehensive Cancer Center grant P30 CA016058. The funders had no role in study design, data collection and analysis, decision to publish, or preparation of the manuscript. The content is solely the responsibility of the authors and does not necessarily represent the official views of the NIH.

## AUTHOR CONTRIBUTIONS

J.K., M.R.K., and G.K., conceptualized this study. J.K., M.R.K., and K.S. performed the experiments. Formal analysis was done by J.K, A.K., and C.T. Data visualization was completed by J.K. Supervision and funding acquisition was done by G.C.K. The initial manuscript and figures were drafted by J.K. and G.C.K. All authors reviewed and edited the final manuscript.

## ETHICS DECLARATIONS

All authors declare no competing interests.

---

**Supplemental Figure 1:**
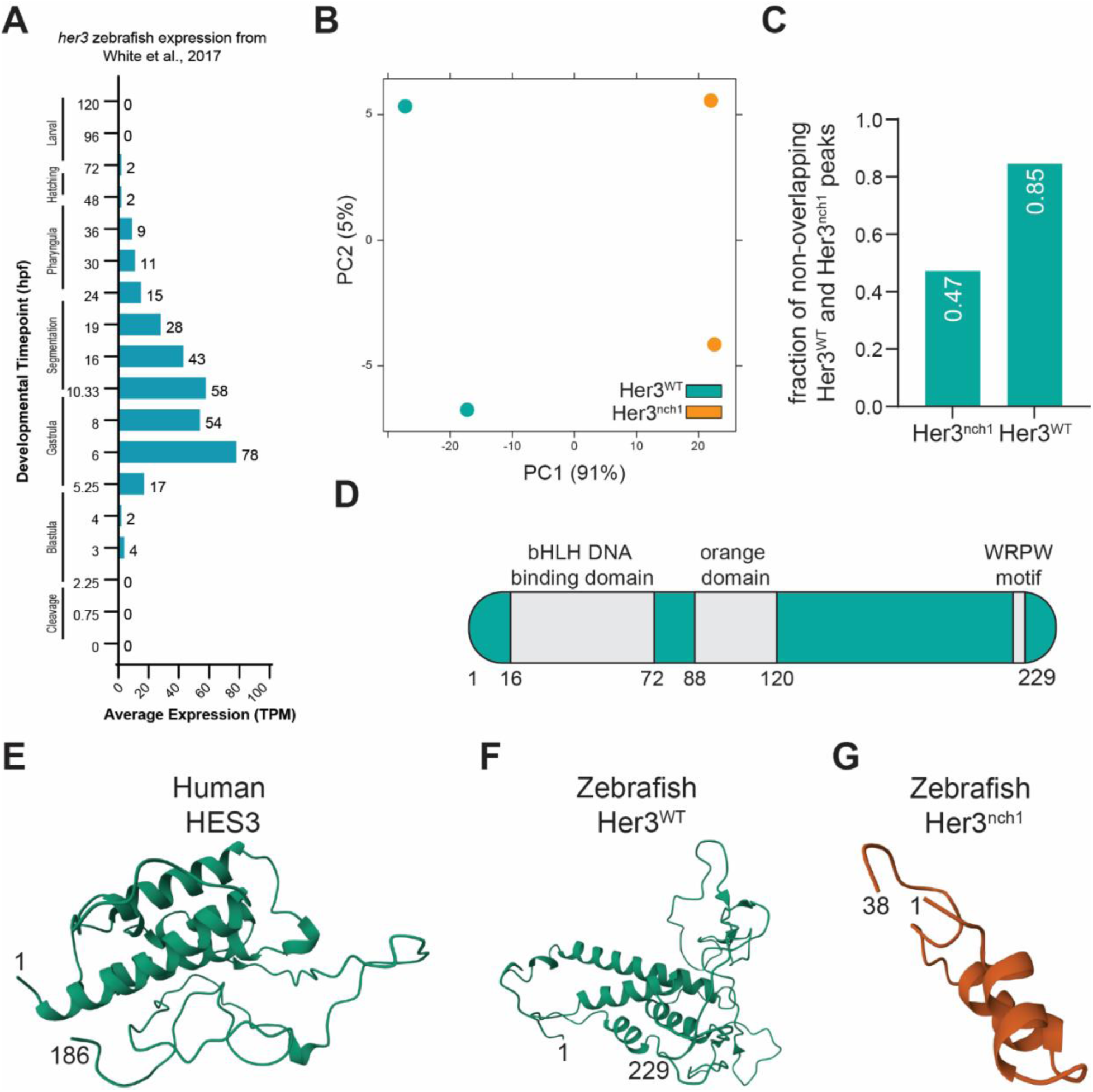
Her3^nch1^ has reduced capability to bind to DNA. (A) *her3* expression during zebrafish embryogenesis from bulk RNA-seq from White et al., 2017.^41^ (B) Principal component analysis on Her3^WT^ and Her3^nch1^ binding sites. (C) Overlap of Her3^WT^ and Her3^nch1^ binding sites with the opposite peak set. (D) Schematic of known wild-type zebrafish Her3 protein domains. (E-G) Structures from molecular dynamics simulations of human HES3, left, zebrafish wild-type Her3, middle, and zebrafish mutant Her3, right.

**Supplemental Figure 2:**
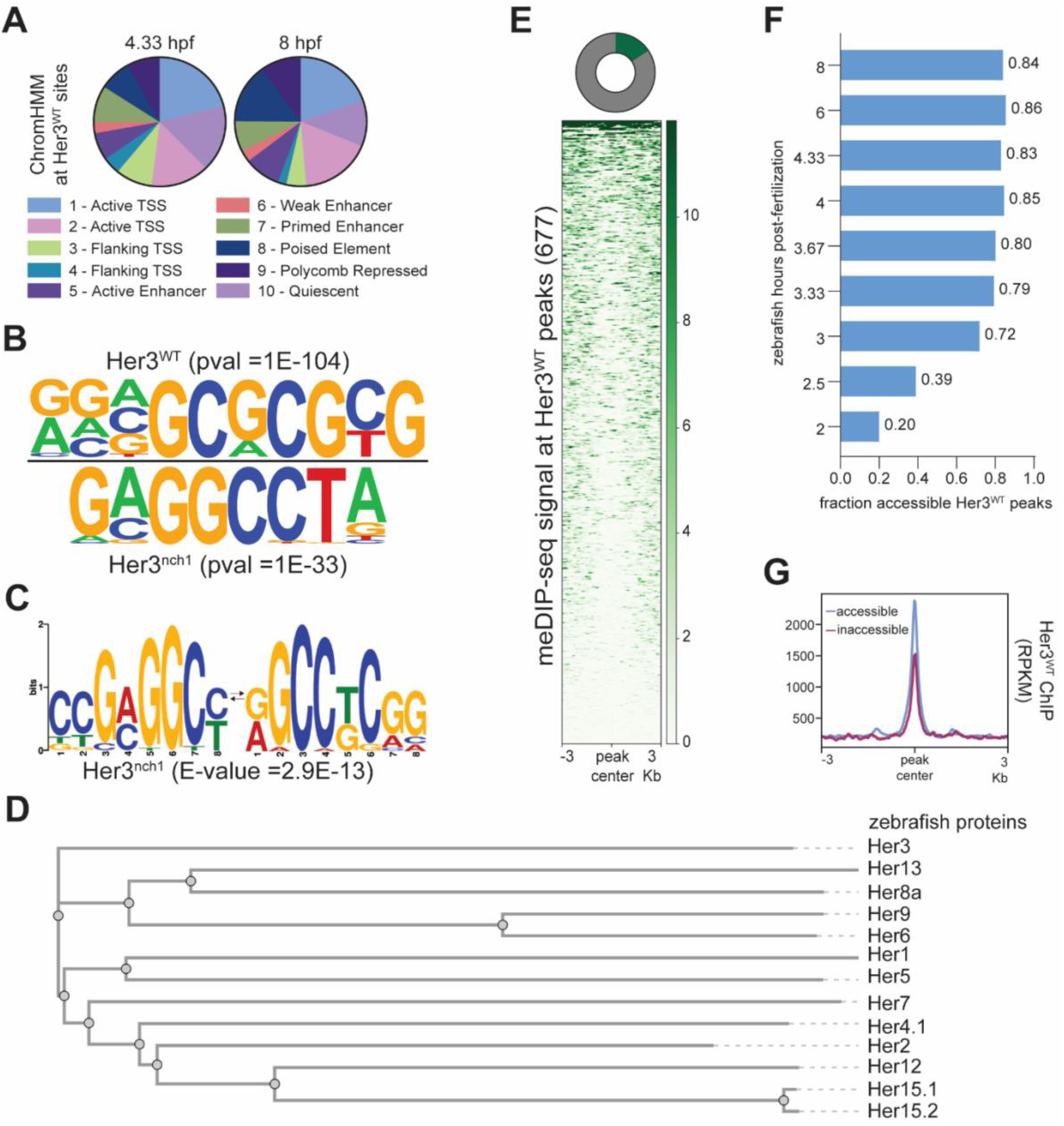
Her3/HES3 binds to open, unmethylated DNA. (A) ChromHMM states at Her3^WT^ binding sites from DANIO-CODE.^90^ (B) Most enriched HOMER de novo motif at Her3^WT^, top, and Her3^nch1^, bottom, binding sites. (C) Most enriched MEME motif at Her3^nch1^ binding sites. (D) Phylogenetic comparison of zebrafish Her (HES) family protein sequences with Clustal Omega.^91^ (E) Top - Overlap between Her3^WT^ binding sites, and meDIP-seq peaks where green represents overlap. Bottom - Heatmap of meDIP-seq signal at Her3^WT^ binding sites. Publicly available meDIP-seq data was obtained from Lee et al., 2015.^45^ (F) Overlap between Her3^WT^ binding sites and ATAC-seq peaks throughout early zebrafish development from publicly available ATAC-seq data from Liu et al. 2018, Pálfy, Schulze et al., 2020., and Kucinski et al., 2025.^39,66,67^ (G) Profile plot of Her3^WT^ ChIP-seq signal at accessible or inaccessible binding sites based on overlap to Kucinski et al., 2025 peaks at six hours post-fertilization.

**Supplemental Figure 3:**
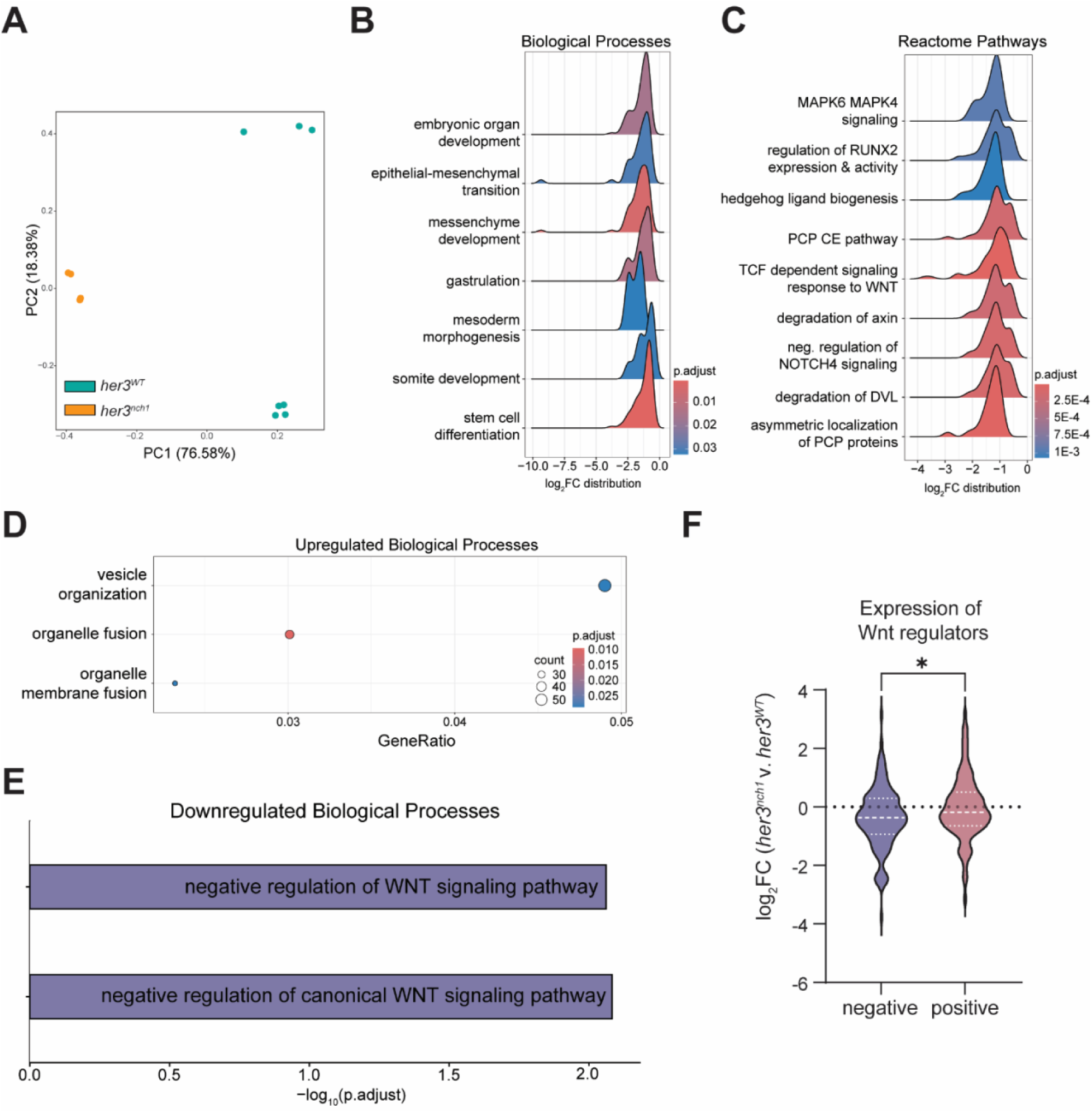
*her3* knockout affects regulators of WNT signaling during zebrafish gastrulation. (A) Principal component analysis on RNA-seq from *her3^nch1^* embryos from this study and Her3 wild-type (*her3^WT^)* embryos from Kucinski et al., 2025.^39^ (B) Ridge plot of gene set enrichment analysis (GSEA) for downregulated biological processes. (C) Ridge plot of GSEA for downregulated Reactome pathways. (D) Over-representation analysis (ORA) of genes upregulated in *her3^nch1^* embryos for biological processes (E) ORA of genes upregulated in *her3^nch1^* embryos for negative WNT regulators. (F) Violin plot of log_2_ fold change (FC) for negative (n=126) or positive regulators (n=110) of WNT signaling. A Mann-Whitney test was used to determine statistical significance. * 0.05 > *p* > 0.01.

**Supplemental Figure 4:**
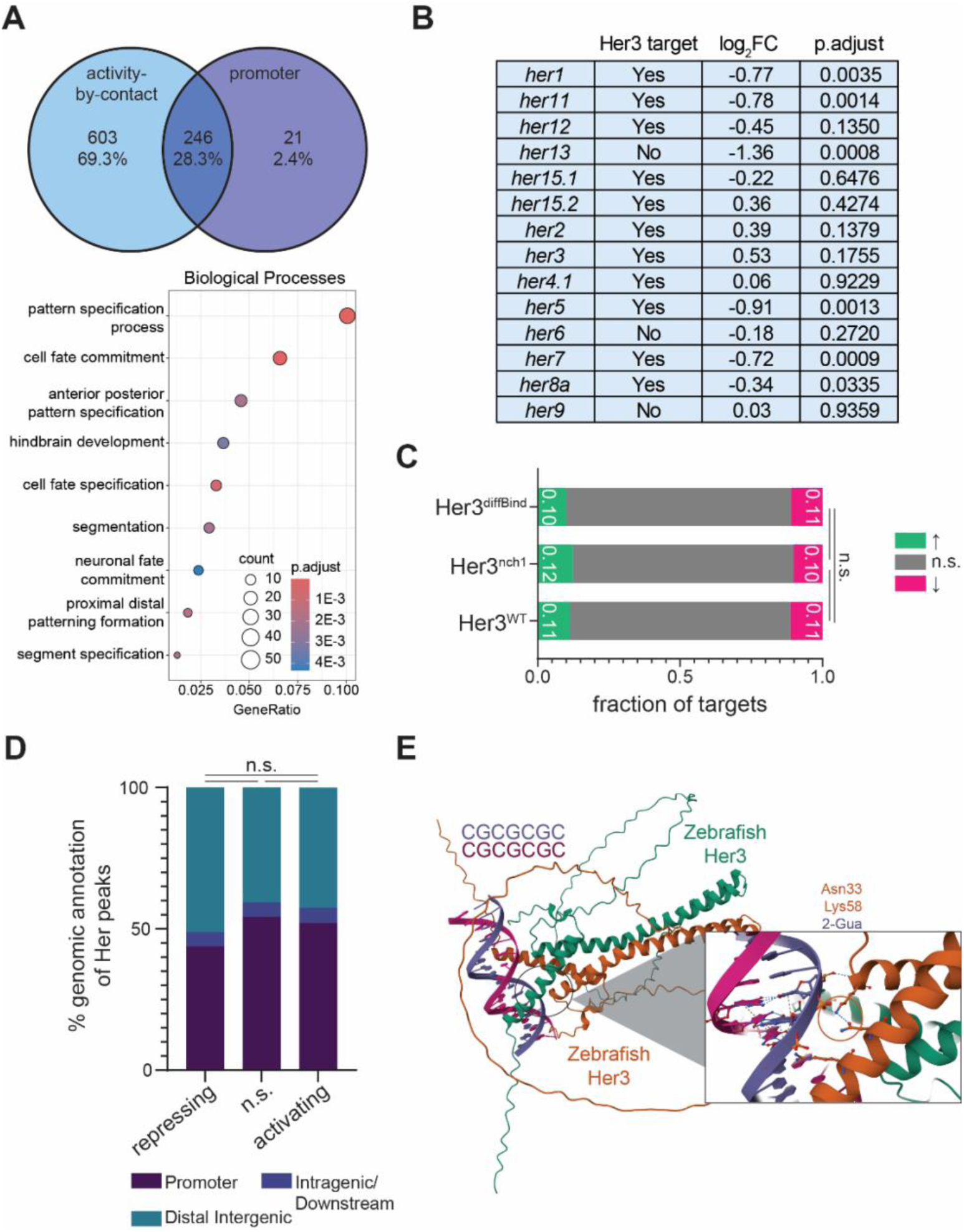
Her3 directly targets developmental and other *her/HES3* genes but does not always induce differential gene expression. (A) Top - Overlap of Her3^WT^ targets that were identified from the activity-by-contact model and promoter binding. Bottom – Over-representation analysis of biological processes for Her3^WT^ target genes. (B) Table showing if *her/HES3* genes are Her3^WT^ targets, left, and if those genes are differentially expressed, middle and right. (C) Fraction of Her3^WT^ targets that are differentially expressed. One-Way ANOVA with Friedman test for multiple comparisons was used to determine statistical significance. (D) Genomic annotation of Her3^WT^ binding sites based on their effect on gene expression. A Fisher’s exact test with a Bonferroni correction for multiple comparisons on the fraction of peaks with promoter binding was used to determine statistical significance. (E) AlphaFold modeling of Her3 binding to the motif at downregulated/activating sites from Figure 2F. n.s. = not statistically significant.

**Supplemental Figure 5:**
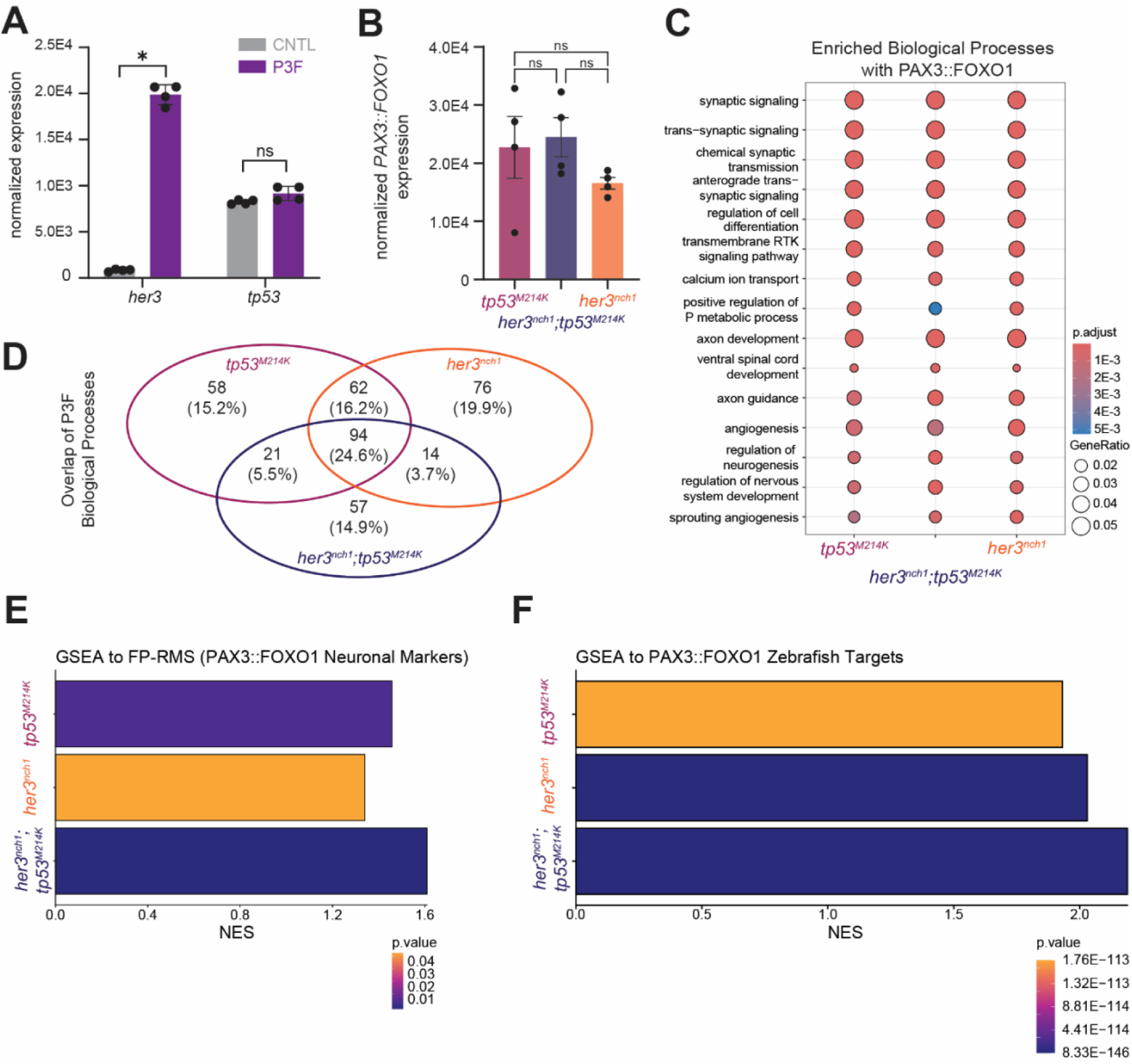
PAX3::FOXO1-injection results in an upregulation of neural programs in *her3* and *tp53* mutant zebrafish embryos. (A) Expression of *her3* and *tp53* after PAX3::FOXO1 mRNA injection in wild-type embryos from Kucinski et al., 2025.^39^ A Mann-Whitney test was used to determine statistical significance. (B) Normalized *PAX3::FOXO1* expression in different mutant zebrafish backgrounds. A Kruskal-Wallis test with a Dunn’s correction for multiple comparisons was used to determine statistical significance. (C) Over-representation analysis of upregulated genes upon PAX3::FOXO1 mRNA injection in different mutant zebrafish backgrounds. (D) Overlap of all upregulated pathways from Supplemental Figure 5C, pathway adjusted *p* < 0.01. (E) Gene set enrichment analysis (GSEA) to PAX3::FOXO1-driven rhabdomyosarcoma neuronal cell population markers identified from scRNA-seq analysis in Danielli, Wei et al. 2023.^32^ (F) GSEA to PAX3::FOXO1 zebrafish targets from Kucinski et al., 2025.^39^ * 0.05 > *p* > 0.01 and n.s. = not statistically significant.

**Supplemental Figure 6:**
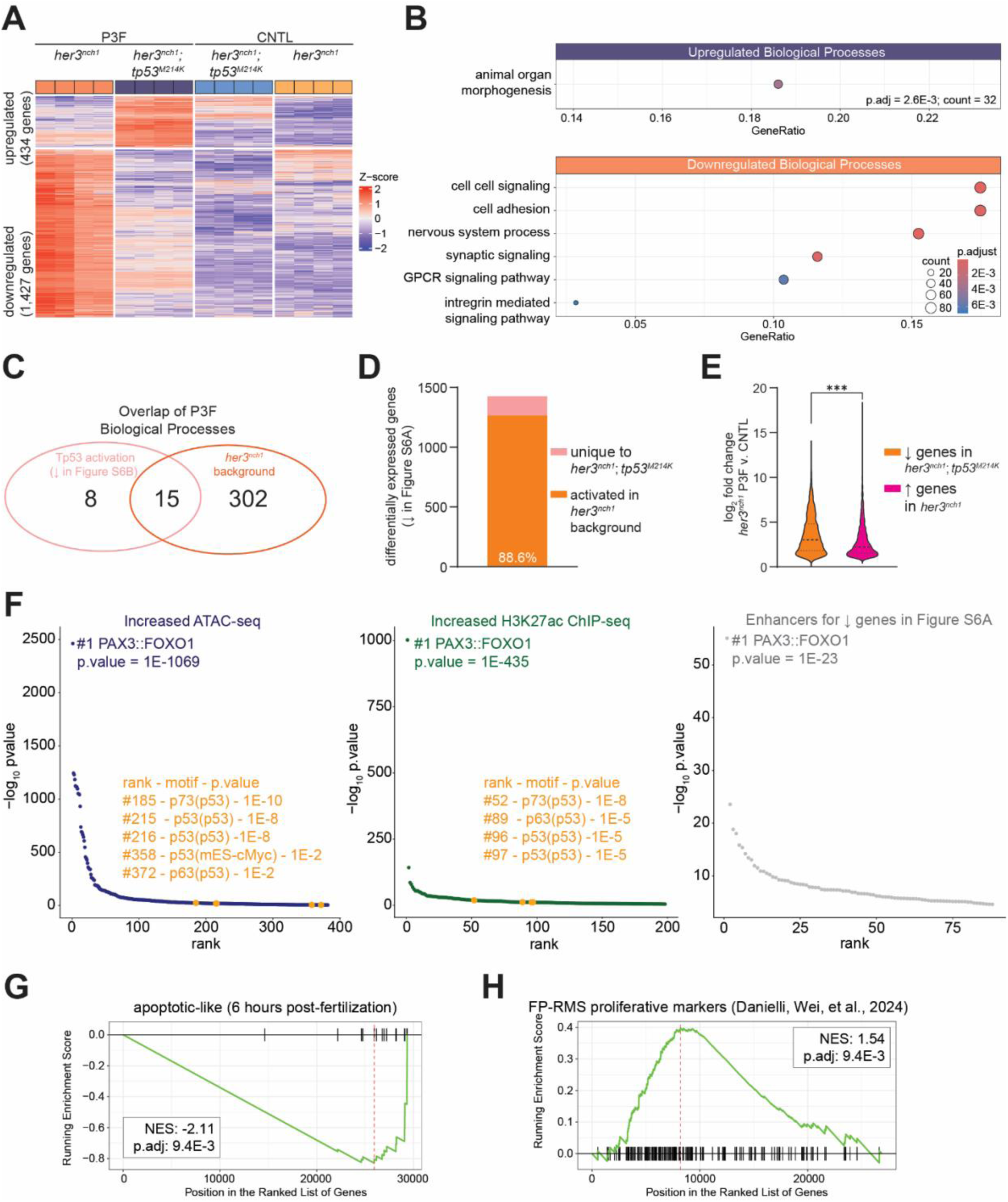
PAX3::FOXO1 in *tp53^M214K^* mutant zebrafish embryos drives further PAX3::FOXO1 gene activation and suppresses apoptotic signatures. (A) Z-score normalized heatmap of differentially activated genes by PAX3::FOXO1 (P3F) in either *her3^nch1^* or *her3^nch1^;tp53^M214K^*embryos. Differentially activated genes from control (CNTL) backgrounds had an absolute fold change ≥ 2.0 and adjusted *p* ≤ 0.05. (B) Over-representation analysis of differentially expressed genes from Supplemental Figure 6A to biological processes upregulated after PAX3::FOXO1 injection in either *tp53^M214K^* or *her3^nch1^;tp53^M214K^*embryos. (C) Overlap of all upregulated pathways from Supplemental Figure 6B, pathway adjusted *p* < 0.01. (D) Number of differentially upregulated genes after P3F-injection in *her3^nch1^;tp53^M214K^*that were already upregulated by P3F in the *her3^nch1^* control background. (E) Violin plot of log_2_ fold change for upregulated genes affected by *tp53^M214K^* in orange versus other upregulated genes upon P3F injection in the *her3^nch1^* control background. A Mann-Whitney test was used to determine statistical significance. \*\*\**p* < 0.001. (F) Relative enrichment of TP53-related motifs at sites with increased accessibility (blue), increased H3K27ac (green), and enhancers for downregulated genes from Supplemental Figure 6A (gray). Peak sets were identified in Kucinski et al., 2025.^39^ (G) Gene set enrichment analysis (GSEA) to markers of the developing zebrafish cell type markers, identified from scRNA-seq analysis from Wagner et al., 2018.^89^ Comparison was made between P3F-injected in *her3^nch1^;tp53^M214K^*versus *her3^nch1^* embryos. (H) GSEA to PAX3::FOXO1-driven rhabdomyosarcoma proliferative cell population markers identified from scRNA-seq analysis in Danielli, Wei et al. 2023.^32^

